# The trait specific timing of accelerated genomic change in the human lineage

**DOI:** 10.1101/2022.02.28.482389

**Authors:** Eucharist Kun, Mashaal Sohail, Vagheesh M. Narasimhan

**Affiliations:** Department of Integrative Biology, The University of Texas at Austin; Centro de Ciencias Genómicas (CCG), Universidad Nacional Autónoma de México (UNAM); Department of Statistics and Data Science, The University of Texas at Austin

## Abstract

Humans exhibit distinct characteristics compared to our primate and ancient hominin ancestors including bipedal locomotion and enhanced neurocognitive ability, but the timing of accelerated changes in these traits is uncertain. To investigate if specific trait-associated variation show enrichment during particular periods of human evolution, we combine genome wide association study (GWAS) data from 70 traits, spanning multiple categories including AI-based image-derived morphological phenotypes of the brain, heart, and skeletal tissues with data from 12 different evolutionary regions obtained from comparative functional genomics, multi-species alignments from long read sequencing, and ancient DNA reflecting 4 different major evolutionary divergence points. These regions cover epigenetic differences in the brain between humans and rhesus macaques, various human accelerated regions (HARs) including regions from the Zoonomia Project, ancient selective sweeps, and Neanderthal introgressed alleles. Using two complementary approaches to examine enrichment between GWAS loci and genomic regions, we show that more phenotypes are enriched in earlier periods of divergence of humans with macaques and chimps, and less so during the divergence with Neanderthals. These traits span respiratory, dermatological, reproductive, metabolic, and psychiatric domains along with skeletal and brain imaging traits, consistent with striking morphological changes between humans and other primates. Among brain imaging traits, we observe an enrichment of SNPs associated with the longitudinal fasciculus in human-gained epigenetic elements since macaques, the visual cortex in HARs, and the thalamus proper in Neanderthal introgressed alleles, implying associated functions such as language processing, decision making, relay of sensory signals, and motor control are enriched at different evolutionary depths.

## Introduction

Humans have distinct features compared to other primates despite our high genetic similarities, encompassing modified skeletal physiology, unique brain size and organization, as well as specific diseases and behaviors^1–5^. Evidence in both the fossil record and comparative anatomy of hominin species reveals shifts in bone structure and size, influencing both our walking patterns and cognitive abilities. In particular, changes in arm length relative to leg length and pelvic morphology facilitates bipedal locomotion, and the increased cranial size in humans results in a brain surface area three times larger than that of our closest living ancestor, the chimpanzee^4,6,7^. Compared to our most recent known hominin ancestors, Neanderthals and Denisovans, there are also variations in limb proportions despite both species also being bipedal, as well as differences in hindbrain and forebrain volume^8^.

While these changes can be observed by examining the fossil evidence and comparisons with the other great apes, it is unclear when along the past few million years that accelerated genomic change for those phenotypes occurred. Two types of datasets from different domains of genomic science may help to address this question. First, over the past decade, there has been major progress in mapping human genotypes to phenotypes. Recent studies have also combined large-scale medical imaging data and genomics to uncover the genetic architecture of skeletal proportions, brain shape and size, as well as heart structure, while biobank data along with large disease-specific cohorts have found novel gene-trait associations across a variety of diseases^9–12^.

Second, on the other side of genomic research, there have been remarkable strides in acquiring and sequencing DNA from both existing and extinct species. New techniques for sequencing ancient DNA have led to the identification of Neanderthal-introgressed alleles, which has unveiled the varying presence of extant hominin DNA in the genomes of modern humans stemming from admixture events around 50,000 to 60,000 years ago^13^. Moreover, advances in sequencing technology, especially long-read sequencing, have broadened the field of comparative genomics, enabling us to generate highly accurate sequences of various organisms^14–17^. By comparing genomic sequences across multiple species, researchers have discovered regions that have acquired more mutations specifically in humans compared with chimpanzees and other mammals^18^. Apart from examining changes in genomic sequences, enhancer elements unique to the human lineage were identified by comparing post translational histone modification profiles across cortical brain tissue in humans, macaques, and mice. These elements, known as human-gained enhancers and promoters (HGEPs), offer a potential avenue for insights into the evolutionary dynamics of the brain over the last approximately 25 million years^19,20^. Through combining genome-wide association studies (GWAS) with evolutionary annotations that have identified human-gained epigenetic marks, accelerated evolution, selective sweeps, and archaic introgression, we can tackle the questions of when human traits likely underwent accelerated evolution.

Previous attempts to answer these questions have used stratified linkage disequilibrium score (S-LDSC) regression to examine the distribution of heritability for phenotypes across the genome. Hujoel et al showed an enrichment of heritability of disease traits in ancient enhancers and promoters as well as in promoters of constrained genes for numerous traits^21^. Other studies have also found that variants introgressed from Neanderthals and shared across multiple Neanderthal populations are enriched in heritability for traits such as skin pigmentation and hair^22–24^.

Lastly, another study discovered heritability enrichment in fetal brain human-gained enhancers as well as Neanderthal introgressed variants (NIVs) for brain MRI phenotypes related to cortical surface area and white matter^25^. These approaches have reliably united GWAS and genomic data to further our understanding of the overall evolution of human traits. We expand on this field of work by using genetic variants associated with 70 independent complex traits, 12 genomic annotations marking sequences that have evolved at different periods of human evolutionary history, and 2 methods of enrichment analysis: S-LDSC as well as a new approach, a gene enrichment pipeline called HARE, which was developed to provide a flexible pipeline for estimating gene overlap enrichment of GWAS traits with genomic regions of interest and provide a complimentary approach to S-LDSC which has known biases in accurately estimating partitioned heritability on smaller size annotations which do not cover enough common SNPs^26,27^.

## Results

### Stratified linkage disequilibrium score regression and gene enrichment analysis of 70 GWAS traits across 12 annotations

We utilized two methods to analyze the enrichment of GWAS trait loci in genomic regions. The first method, stratified LD score regression (S-LDSC), partitions heritability from GWAS summary statistics based on genomic regions, known as annotations, and determines if a particular region explains more heritability for a given trait than would be expected from the proportion of SNPs present in that annotation^28^. For this method, we analyzed our test annotations in a model simultaneously incorporating several other regulatory elements, measures of selective constraint, and linkage statistics (baselineLDv2.2 with 97 annotations) to estimate heritability enrichment (*h*^2^(*C*)) while minimizing bias due to model misspecification (**Methods**: Stratified LD score regression (S-LDSC) framework)^21,28–30^. The second method, HARE, maps genome-wide significant SNPs to nearby genes and scans for elevated levels of overlap between these genes and genomic annotations of interest. We then estimate overlap enrichment (*I*) by calculating the percent difference between the amount of overlap between a test annotation and a set of genes related to a GWAS trait versus the amount of overlap between the annotation and a random set of genes matched for gene length and number of genes for each trait (**Methods**: Gene enrichment analysis for evolutionary annotations)^27^. Percentage difference values greater than 0% are considered enriched in an annotation.

In order to examine a range of complex traits and diseases that cover human specific phenotypes as well as those common to other great apes, we chose a set of 42 relatively unrelated GWAS datasets previous analyzed in Hujoel et al. across distinct trait domains including psychiatric, immunological, dermatological, skeletal, reproductive, metabolic, cardiovascular, gastrointestinal, or endocrine as defined in the GWAS Atlas (heritability (hg) = 0.92% -0.67%, genetic correlations (rg) < 0.9, average sample size (N)=320,000)^21,31^. Additionally, we analyzed GWAS results from 3 image-derived phenotyping studies carried out on medical images. These include 16 brain MRI phenotypes spanning brain size, structure, and MRI activity, 6 right heart MRI phenotypes related to atrium size and function, and 6 skeletal dual-energy X-ray absorptiometry (DXA) phenotypes related to skeletal proportions (hg = 8% -51%, rg < 0.9, N = 31,221 – 41135) (**Fig. 1**, **Table S1** and **Table S2**)^11,32,33^.

**Fig. 1.**
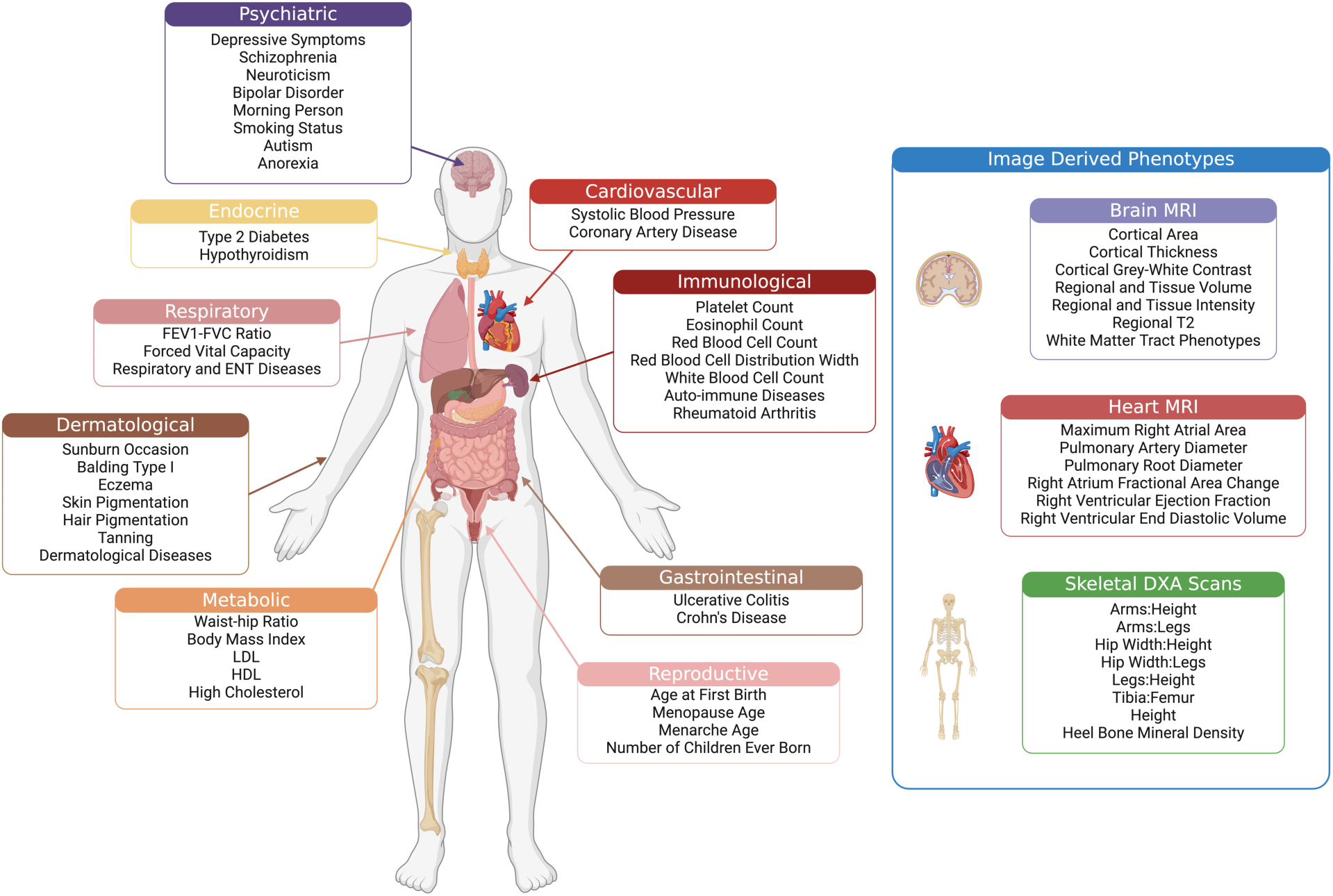
Phenotypes examined in our gene and heritability enrichment analyses. Phenotypes were grouped according to study and/or GWAS Atlas category (psychiatric, endocrine, respiratory, dermatological, metabolic, cardiovascular, immunological, gastrointestinal, and reproductive). Morphological phenotypes of the human body obtained specifically from GWAS carried out on UK Biobank medical imaging cohorts (brain MRI, heart MRI, and full body DXA) are shown to the right and grouped in a blue box.

We chose to examine the role of comparative functional genomic data, multispecies alignments from long read sequencing, and ancient DNA in shaping polygenic traits and disease evolution across a timescale of 25 million years through these genomic annotations marking various evolutionary periods: (A) Human-gained enhancers and promoters (HGEPs) in the brain since human divergence with rhesus macaque at different post-conception weeks (p.c.w) and adulthood^19,20^, (B) The fastest evolving unique regions of the human genome when compared to various mammal and primate genomes (human accelerated regions (HARs), lineage-specific accelerated regions (LinARs), high-confidence HARs (zooHARs), and human ancestor quickly evolved regions (HAQERs))^16,17,34,35^, (C) Ancient selective sweeps in humans relative to Neanderthals and Denisovans^36^, and (D) Neanderthal introgressed variants^37^. These genomic annotations constitute different numbers of regions, genomic sizes, and proportion of total common variation (**Fig. 2**, **Table S3**). Because of previously mentioned S-LDSC limitations for accurately estimating partitioned heritability on smaller size annotations as well as strength to control for epigenetic annotations in different cell types, we performed S-LDSC analysis on only the HGEPs and HARE analysis on the remaining evolutionary annotations^26^.

**Fig. 2.**
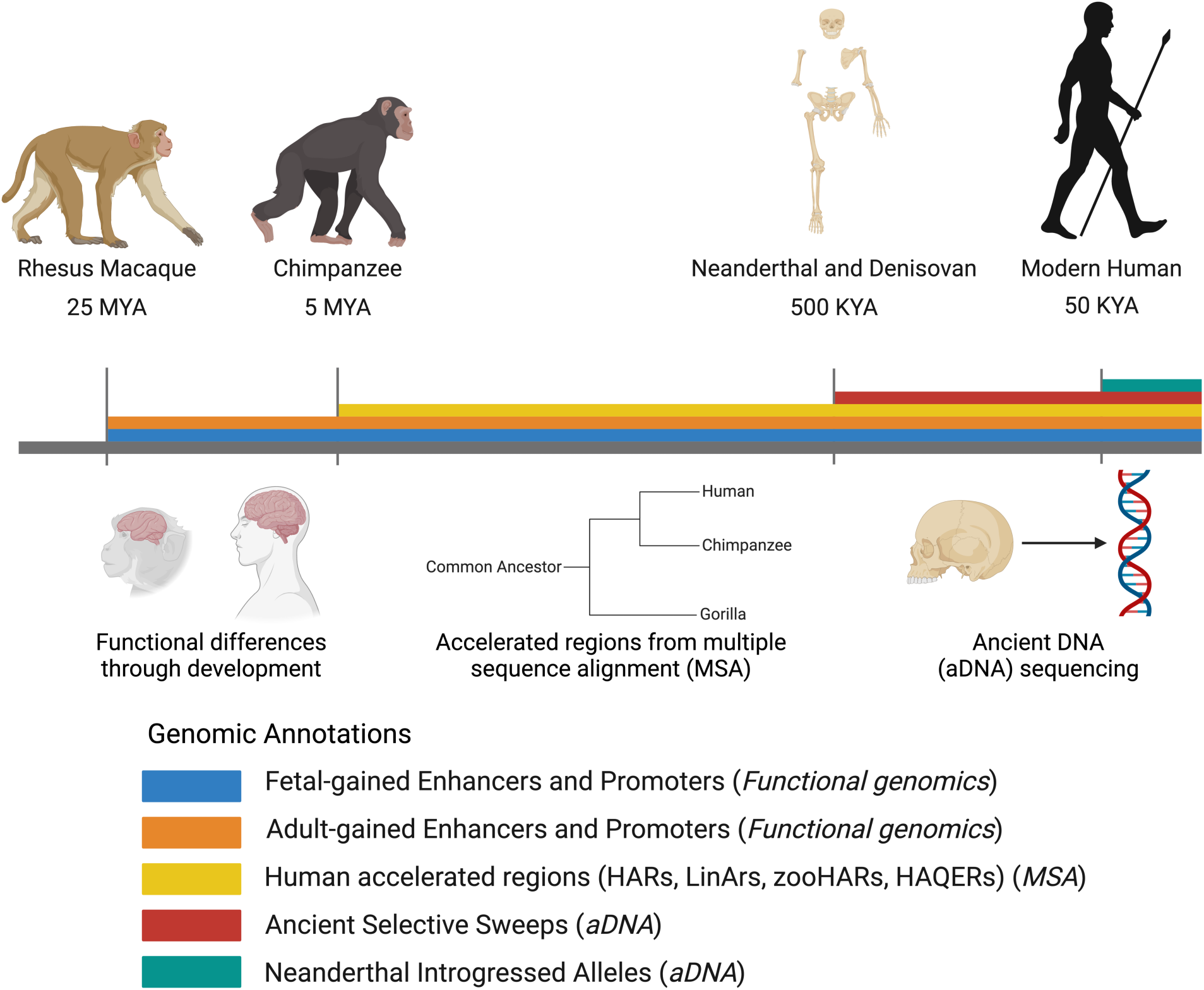
Evolutionary time periods. Major time points of primate evolution relevant in this study are highlighted. Genomic annotations corresponding to evolutionary time periods are shown in color on the timeline. These annotations include fetal HGEPs (blue), adult HGEPs (orange), human accelerated regions (yellow), ancient selective sweeps (Extended Lineage Sorting ELS) (red), and putatively introgressed variants from Neanderthals (teal). Methods used to obtain each genomic region of interest are also shown below their corresponding annotations. Blue and orange intervals mark epigenetic gains in the cerebral cortex while the other color intervals mark genetic gains discovered through comparative genomics from primate and ancient DNA sequences.

### Fetal human-gained enhancers and promoters in early development are enriched in traits across skeletal, respiratory, and dermatological domains

First, considering the period from our divergence from our most recent common ancestor with rhesus macaque (∼25 million years ago (MYA)), we analyzed HGEPs in the brain at 7, 8, and 12 p.c.w and adulthood for a total of 6 annotations. We performed S-LDSC on 70 traits across the 6 annotations while performing additional control by adding all epigenetic fetal and adult brain regulatory elements from the Epigenome Roadmap Project 25 state model to our model (**Methods**: Stratified LD score regression (S-LDSC) framework). We found that fetal human-gained regulatory elements at 7 p.c.w were enriched in heritability for traits across several domains at FDR-adjusted p-value < 0.05 (heel T score, balding type I, FEV1-FVC ratio, height, forced vital capacity, systolic blood pressure, skeletal proportions, and weighted-mean isotropic or free water volume fraction (ISOVF) in tract left superior longitudinal fasciculus) (*h*^2^(*C*) = 4.26 – 11.30, p = 4.00 × 10^-2^ – 6.94 × 10^-5^) (**Fig. 3**). Comparatively, HGEPs at other time points either fetal or adult showed little to no enrichment for specific traits, with only forced vital capacity showing significant enrichment at 8 p.c.w (*h*^2^*C*) = 6.52, p = 3.84 × 10^-2^) (**Table S4**).

**Fig. 3.**
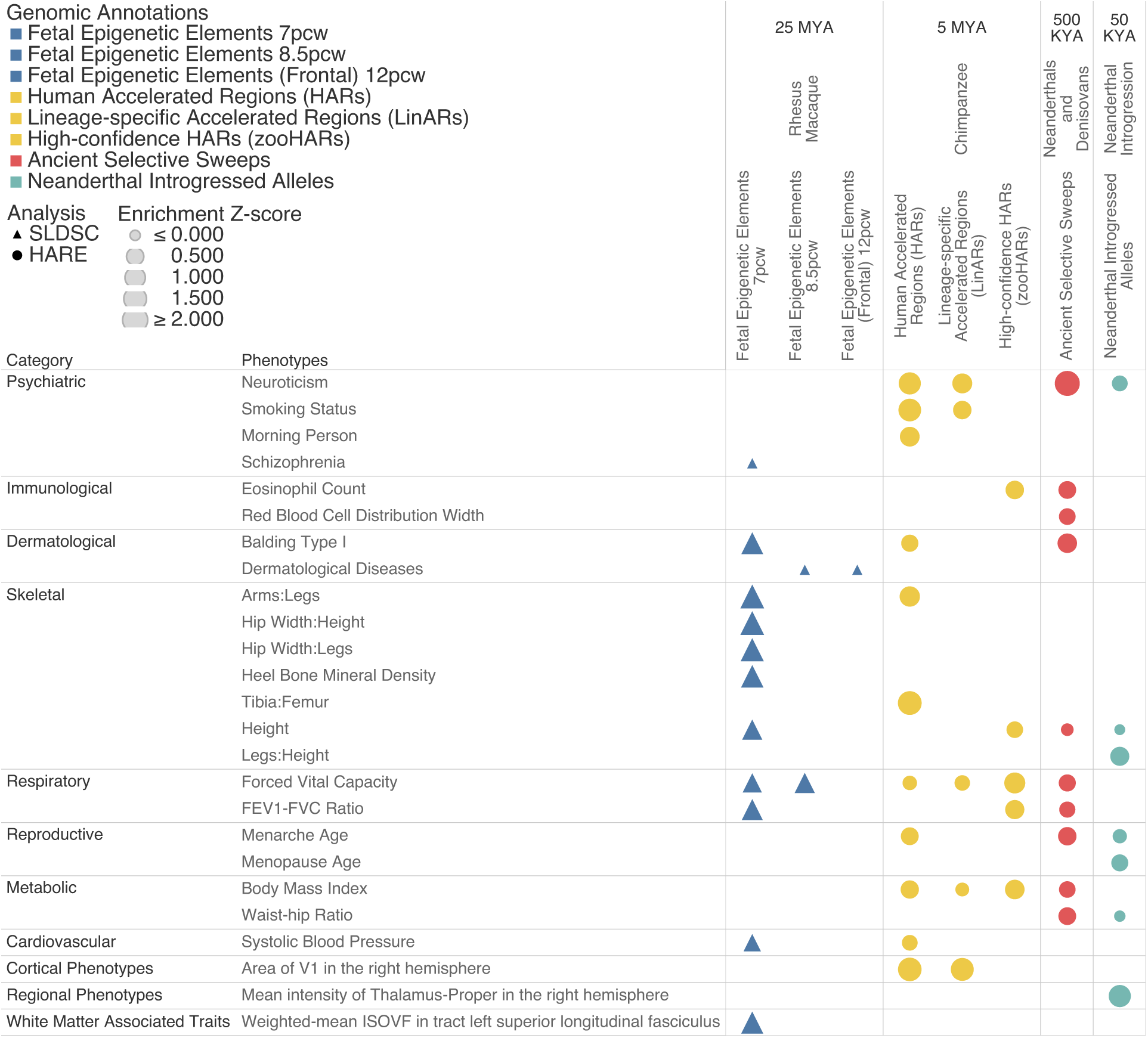
Heritability and gene enrichment of human phenotypes across evolutionary time periods. S-LDSC and HARE analysis of 70 traits for 12 genomic annotations marking various time periods in human evolutionary history. Color corresponds to each major evolutionary period analyzed in accordance with the schema in **Fig. S2** and each shape marks the type of analysis performed - triangular for S-LDSC and circular for HARE. Enrichment z-scores were calculated separately for each type of analysis and the resulting enrichment value is denoted by the overall size of each shape. All results shown are significant at FDR-adjusted p-value < 0.05.

To check whether our heritability enrichment was affected by tissue specificity, we performed a secondary S-LDSC analysis on the 7 p.c.w time point annotation. We obtained genomic regions pertaining to regulatory elements from various tissues in the Epigenome Roadmap 25 state model and created an annotation set of enhancers and promoters that were common to all major tissues in the Roadmap Epigenome Consortium dataset^38^. We intersected the human gained elements found specifically in the brain with this pan tissue annotation set to generate two subsets of annotations, regulatory elements that overlapped with genomic regions shared by Epigenome Roadmap tissues and elements that had no overlap. We then performed S-LDSC on each set and correlated the heritability enrichment values of each subset with the original annotation set. In doing so, we found that our heritability enrichment results for both subsets were over 90% correlated with the original annotation, and we confirmed that HGEPs found only in brain tissue at 7 p.c.w contribute significant heritability enrichment across various domains including dermatological, skeletal, and respiratory traits (*h*^2^(*C*) 4.53 – 12.55, p = 9.56 × 10^-3^ – 1.23 × 10^-4^) (**Fig. S1**) (**Table S5**) (**Methods**: Tissue specificity analysis). This suggests that the majority of the evolutionary enrichment signals we observe are driven by a core set of enhancers and promoters that are tissue agnostic that are differentially regulated in humans vs rhesus macaques, and is also suggestive of pleiotropic effects across the brain-body axis through epigenetic regulation.

### Various sets of human accelerated regions are enriched for psychiatric, respiratory, and metabolic traits as well as the area of the visual cortex

We then carried out gene enrichment analysis on various types of human accelerated regions of the genome, HAQERs, HARs, LinARs, and zooHARs, which all pertain to the period spanning the last known common ancestor between humans and chimpanzees (∼5 MYA). These comparative genomic annotations were generated through different genomic species alignments and methodologies and share various amounts of overlap as 18% of LinARs overlap with HARs and 45% of zooHARs overlap with HARs, but each annotation also exhibits distinct features (**Methods**: Differences among various types of human accelerated regions). Our results show that enrichment analysis for HARs and LinARs were largely similar (Pearson correlation = 0.71), with any trait that was significantly enriched in LinARs also being significantly enriched in HARs (neuroticism, smoking status, forced vital capacity, BMI, and area of the primary visual cortex in the right hemisphere). However, zooHARs deviated slightly from HARs (Pearson correlation = 0.42) and LinARs (Pearson correlation = 0.29) and had unique significance (FDR-adjusted p-value < 0.05) in several traits (eosinophil count, height, and FEV1-FVC ratio) (*I* = 68%, p = 3.47 × 10^-2^; *I* = 51%, p = 9.52 × 10^-5^; *I* = 76%, p = 1.30 × 10^-2^) (**Fig. 3**). BMI and forced vital capacity were significantly enriched across these three types of accelerated regions (*I* = 37% – 100%, p = 2.36 × 10^-3^ – 2.18 × 10^-4^; *I* = 34% – 81%, p = 3.27 × 10^-4^ – 4.84 × 10^-84^) and neuroticism, smoking status, and area of the primary visual cortex in the right hemisphere were enriched across HARs and LinARs (*I* = 85% – 117%, p = 3.72 × 10^-4^ – 8.78 × 10^-23^; *I* = 68% – 126%, p = 4.63 × 10^-3^ – 1.96 × 10^-27^; *I* = 130% – 141%, p = 2.67 × 10^-2^ – 1.14 × 10^-3^) (**Fig. 3**) (**Table S6**). As HAQERs are found in the non-conserved genome and do not typically overlap any genes, there was no enrichment for any trait which falls in line with our expectations from this type of analysis (**Methods**: Differences among various types of human accelerated regions)^35^.

### Ancient selective sweeps and Neanderthal introgressed regions are enriched in immunological, dermatological, respiratory, and reproductive traits

Lastly, we examined human evolution over the past several hundred thousand years by carrying out gene enrichment analysis on ancient selective sweeps and Neanderthal introgressed regions. We observed significant enrichment (FDR-adjusted p-value < 0.05) for several traits and trait domains in both types of annotations. In agreement with hypotheses regarding differences between Neanderthals and humans, immunological, dermatological, and respiratory traits were enriched in ancient selective sweeps (eosinophil count, red blood cell distribution width, balding, FEV1-FVC ratio, and forced vital capacity) (*I* = 47% – 80%, p = 2.03 × 10^-2^ – 8.83 × 10^-4^) while reproductive traits were enriched in introgressed alleles (menarche and menopause age) (*I* = 36%, p = 2.22 × 10^-4^; *I* = 54%, p = 2.02 × 10^-3^) (**Fig. 3**)^23,39–41^. Of particular interest, our analysis found that genes associated with the thalamus, which is responsible for relaying sensory information in the brain, were also enriched in NIVs (*I* = 117%, p = 1.93 × 10^-2^) as well as certain body proportions known to differ between modern humans and Neanderthals (waist-hip ratio and leg-height ratio) (*I* = 28%, p = 2.65 × 10^-4^; *I* = 74%, p = 2.55 × 10^-2^) (**Fig. 3**) (**Table S6**)^8^.

### Meta-analysis of S-LDSC and gene overlap enrichment show significant enrichment across several domains

To determine if overall trait domains were enriched in genomic annotations pertaining to specific evolutionary time periods, we carried out a meta-analysis across our GWAS categories (**Methods**: S-LDSC and HARE meta-analysis). Overall, the meta-analysis of the S-LDSC results shows that the highest number of significantly enriched categories (FDR-adjusted p-value < 0.05) are in the 7 p.c.w time point (immunological, skeletal, respiratory, gastrointestinal, cortical brain MRI, regional brain MRI, and heart MRI), and that respiratory and skeletal traits are enriched across all fetal timepoints (**Fig. 4**). Additionally, psychiatric and cardiovascular traits are the sole categories to show an increase in enrichment across the developmental periods and only show significant enrichment at the 12 p.c.w time period (*h*^2^(*C*) = 2.56, p = 3.40 × 10^-3^, *h*^2^(*C*) = 3.20, p = 1.09 × 10^-12^) in occipital and frontal regions respectively (**Fig. 4**) (**Table S7**). Lastly, white matter associated traits which form the network of the brain by connecting different brain structures and play a crucial role in cognition, particularly cognitive speed, were the only category to be significantly enriched in heritability in adult human gained enhancers and was not enriched in any other developmental period^42,43^. We performed the same meta-analysis using the HARE results and found only a few categories of traits to be significantly enriched (FDR-adjusted p-value < 0.05) across the genomic annotations. These include psychiatric traits in HARs (*I* = 100%, p = 4.32 × 10^-7^) and dermatological and respiratory traits in zooHARs, regions accelerated in our genomic evolution since divergence with chimps (*I* = 84%, p = 2.13 × 10^-9^; enrichment = 0.92, p = 1.53 × 10^-8^), but only dermatological traits were significantly enriched in regions under selection since our divergence with Neanderthals (*I* = 35%, p = 6.82 × 10^-4^) (**Fig. 4**) (**Table S8**).

**Fig. 4.**
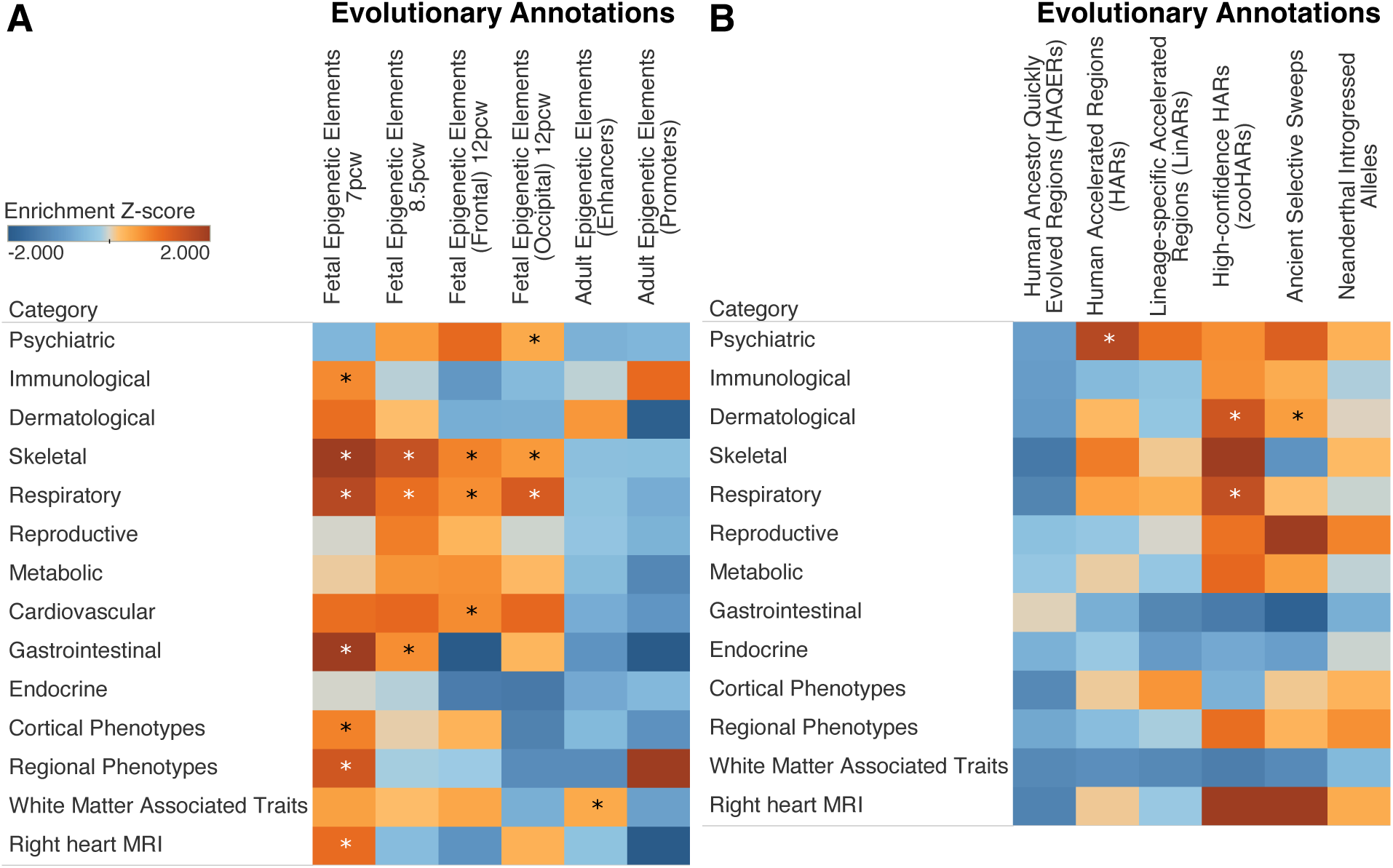
Meta-analysis of heritability and gene enrichment. **A**) Meta-analysis of S-LDSC results across HGEPs based on trait categories. **B**) Meta-analysis of HARE results across evolutionary annotations based on trait categories. Asterisks denote significance at FDR-adjusted p-value < 0.05 across each type of analysis.

### Genomic annotation robustness and tissue overlap sensitivity analysis

As many of the genomic annotations we used in our analysis were derived from computational approaches that may not be able to accurately determine these regions at base pair level resolution, we ran additional sensitivity analyses to test the robustness of both the various methods used to generate the annotations we tested as well as our enrichment methods. We accomplished this by modifying our genomic regions of interest in various ways. First, we examined the impact of randomly sampling just 90% of each of the annotations over 3 replicates and running S-LDSC and HARE on these subsets (**Fig. S2**). Additionally, we took a separate approach where we modified the lengths of each individual region within each annotation, first making each region 10% smaller followed by making each region 10% larger and ran S-LDSC and HARE on each modified subset. We then correlated the corresponding enrichment estimates and p-values to the original results and found that all three types of modified annotation sets had an average Pearson correlation of 95% with the original annotations (**Methods**: Annotation robustness analysis) (**Table S9** to **Table S11**).

Additionally, to determine whether the significant heritability enrichment we observed in fetal gained enhancers and promoter at 7 p.c.w and not the other fetal timepoints was largely due to overlap with epigenetic elements present in other tissues, we examined base pair overlap among all fetal timepoint annotations with regulatory elements from various tissues in the Epigenome Roadmap 25 state model (**Table S12** – **Table S14**)^38^. Our results show that base pair overlap between the brain and other tissues actually increases on average from 7 p.c.w to 12 p.c.w, indicating that the signals we observe in the 7 p.c.w annotation are not solely driven by greater overlap with regulatory regions as compared to the other fetal timepoint annotations (**Fig. S4** and **Fig. S5**) (**Methods**: Tissue specificity analysis).

## Discussion

Overall, our study examines when various phenotypes underwent accelerated evolution across developmental time points and evolutionary events ranging from human divergence from macaques to ancient hominin introgression. We first examined evolutionary changes in the brain since our divergence from rhesus macaque over 25 MYA using S-LDSC and discovered that 12 of our 70 traits showed heritability enrichment in this time period. Across the various developmental time points, the 7 p.c.w period showed heritability enrichment in far more traits and trait domains than other stages of development as determined by both the individual and the meta-analysis. Our results demonstrate that epigenetic elements that gained novel function in the fetal brain since human divergence with rhesus macaque are likely involved in the development of several skeletal, dermatological, and respiratory traits at 7 p.c.w. suggesting pleiotropic effects across the brain-body axis from our early evolution and life history. Next, we analyzed 4 types of fast evolving regions of the human genome since our divergence from chimpanzees over 5 MYA using HARE and found that 14 traits showed significant gene enrichment for these genomic regions, and a meta-analysis of our results showed that psychiatric traits overall were also significantly enriched in HARs. Though different methods and genomic alignments spanning various species were used to generate these human accelerated regions of the genome, our results largely show similar trait enrichment across each annotation (**Methods**: Differences among various types of human accelerated regions). Lastly, we analyzed the last 50 - 100 thousand years of human evolution through leveraging information from Neanderthal and Denisovan DNA and discovered that 10 traits showed gene enrichment in ancient selective sweeps and 7 traits showed gene enrichment in Neanderthal introgressed alleles. In line with observed changes in the fossil record and previous genomic evidence linking selective sweeps and introgressed hominin DNA in humans to various adaptive phenotypes, these traits spanned immunological, dermatological, skeletal, respiratory, reproductive, and metabolic categories. Overall, our results show that more traits were enriched between the periods of divergence related to humans and macaques as well as humans and chimpanzees rather than the period of divergence related to Neanderthals and Denisovans.

Of particular interest, traits that were enriched in multiple annotations showed continuity over consecutive time periods. For example, we observed that BMI was enriched in both HARs and ancient selective sweeps while height and waist to hip ratio were enriched in ancient selective sweeps and NIVs, but no traits were enriched in only HARs and NIVs or only HGEPs and NIVs. Balding and forced vital capacity were enriched across HGEPs at 7 p.c.w, HARs, and ancient selective sweeps while neuroticism and menarche age were significantly enriched across HARs, ancient selective sweeps, and NIVs. Balding, forced vital capacity, and menarche age are related to observed or hypothesized phenotypic differences in lung function, hair, and reproduction across primates, Neanderthals, and humans while neuroticism could be associated with general changes in behavior and emotional complexity which also differ in humans from our evolutionary ancestors, and our study provides firsthand genomic evidence of accelerated evolution for these traits across multiple periods of evolution^23,44–48^.

Across the deep-learning-based image derived phenotypes of the skeleton, brain, and heart, we show evidence that all three major body systems underwent accelerated change in our human lineage. In particular, our meta-analysis results show that all three modalities were enriched in HGEPs since our divergence with macaques, and individual skeletal traits such as arm to leg ratio and leg to height ratio as well as brain traits such as area of the visual cortex and intensity of the thalamus were also enriched in accelerated regions as well as introgressed alleles (**Fig. 3**), hinting at continued evolution of these traits across timepoints. These observations mirror common observed morphological differences between humans and other great apes as our skeleton structures, brain size and organization, and heart morphology have all undergone significant changes^49–52^.

Looking closer at the three brain imaging traits that were enriched at different evolutionary timepoints, we found that each was related to a brain MRI category as well as a substructure of the brain. In our earliest evolutionary timeframe regarding human divergence with macaques, we found that white matter substructure measurements in the left superior longitudinal fasciculus, which is part of the complex network connecting different brain structures to each other and is associated with language processing, decision making, and motor function, were enriched in fetal-gained epigenetic elements at 7 p.c.w^53^. Next, we found that the area of the visual cortex in the right hemisphere, which processes visual information from the retinas, was enriched in human accelerated regions since our divergence with chimpanzees^54^.

Lastly, the mean intensity of the thalamus proper in the right hemisphere, which is primarily associated with relaying information from vision, taste, touch, and hearing sensory pathways to the cerebral cortex as well as relaying motor signals from the cerebellum to the motor cortex, was enriched in Neanderthal introgressed alleles^55^. The temporal approach of this study paints a possible roadmap of the differential timepoints various brain structures underwent accelerated evolution.

Our work as well as other forays into genomic enrichment analysis to take advantage of continuous advances in generating large amounts of GWAS and genomic annotation data. As such, we face similar limitations outlined by Hujoel et al and Alagoz et al^21,25^. Firstly, our data is limited only to GWAS carried out on populations with European ancestry due to a lack of large-scale cohorts of more diverse populations. Additionally, although we use some of the current largest sample size GWAS available, some of the traits we analyze are likely insufficiently powered for truly accurate heritability estimation. Our work would greatly benefit from increases in GWAS data through larger and more diverse sample sizes as well as study of novel traits.

However, the key limitation is one of evolutionary annotation. While ancient DNA has been a revolutionary technology in our ability to obtain genetic data from our archaic ancestors, there is a lack of genomic data between human divergence from chimpanzees and the emergence of Neanderthals. Some of the most overt changes to the human physiological form such as bipedal walking and increased cranial size occurred during this particular period and is evident through the hominin fossil record^56^. An important avenue moving forward with this work would involve advances in aDNA sequencing techniques to elucidate genomic changes that occurred from ∼5 MYA to 500 thousand years ago (KYA) across various extinct hominin species. Even using the latest methods, high quality ancient human genomes are only available for samples up to 40,000 years old and no ancient DNA has ever been extracted from any sample dating older than 430 KYA^57,58^. Lastly, our analysis of HGEPs since divergence with rhesus macaque was limited to epigenetic elements found in the brain tissue. As many of our analyzed phenotypes may not be affected by brain specific HGEPs, our work can only give a partial glimpse at accelerated evolutionary trends during this time period. Further work to discover more human-gained enhancers and promoters across a variety of tissues would provide an opportunity for a more unbiased approach of examining when human traits experienced accelerated evolution.

Overall, in this study we probe a variety of complex traits and diseases for accelerated evolution across four major points of human evolutionary divergence related to macaques, chimpanzees, and Neanderthals/Denisovans ranging from 25 MYA to 50 KYA. We found significant enrichment across various time points for traits spanning from psychiatric-related to skeletal morphology. Combined, we paint a temporal landscape of when phenotypes underwent accelerated evolution in the human lineage.

## Acknowledgments

Figs 1 and 2 were created with Biorender.

## Author contributions

M.S. conceived of the initial project. M.S. and E.K. performed analysis. E.K., M.S. and V.M.N. wrote the paper. M.S. and V.M.N. jointly supervised the work.

## Supplementary Figures

**Fig. S1.**
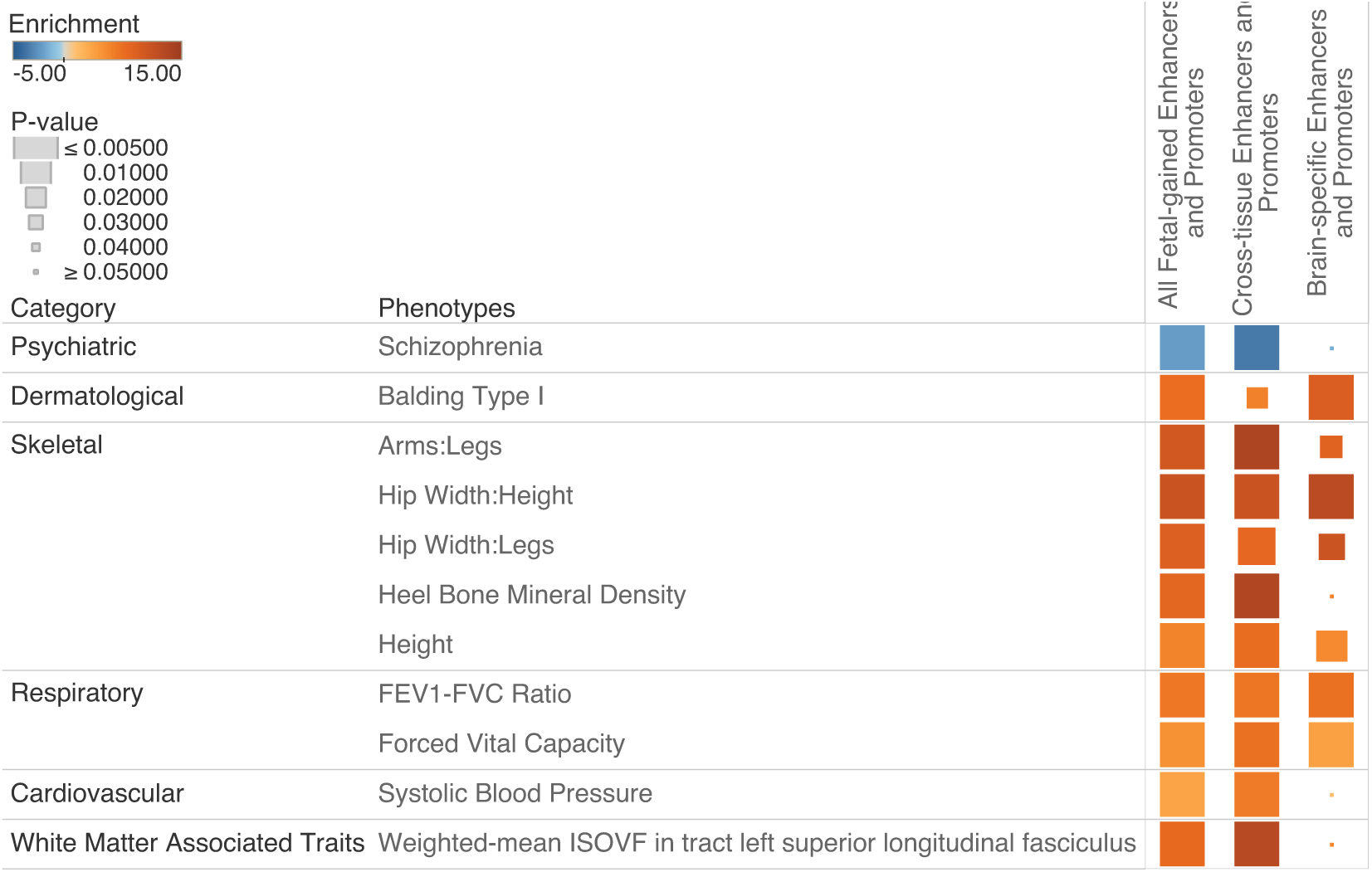
Comparison of S-LDSC heritability enrichment across fetal-gained enhancers and promoters at 7 p.c.w in the brain. Heritability enrichment carried out on the original set shown on the left column followed by cross-tissue enhancers and promoters then brain-specific enhancers

**Fig. S2.**
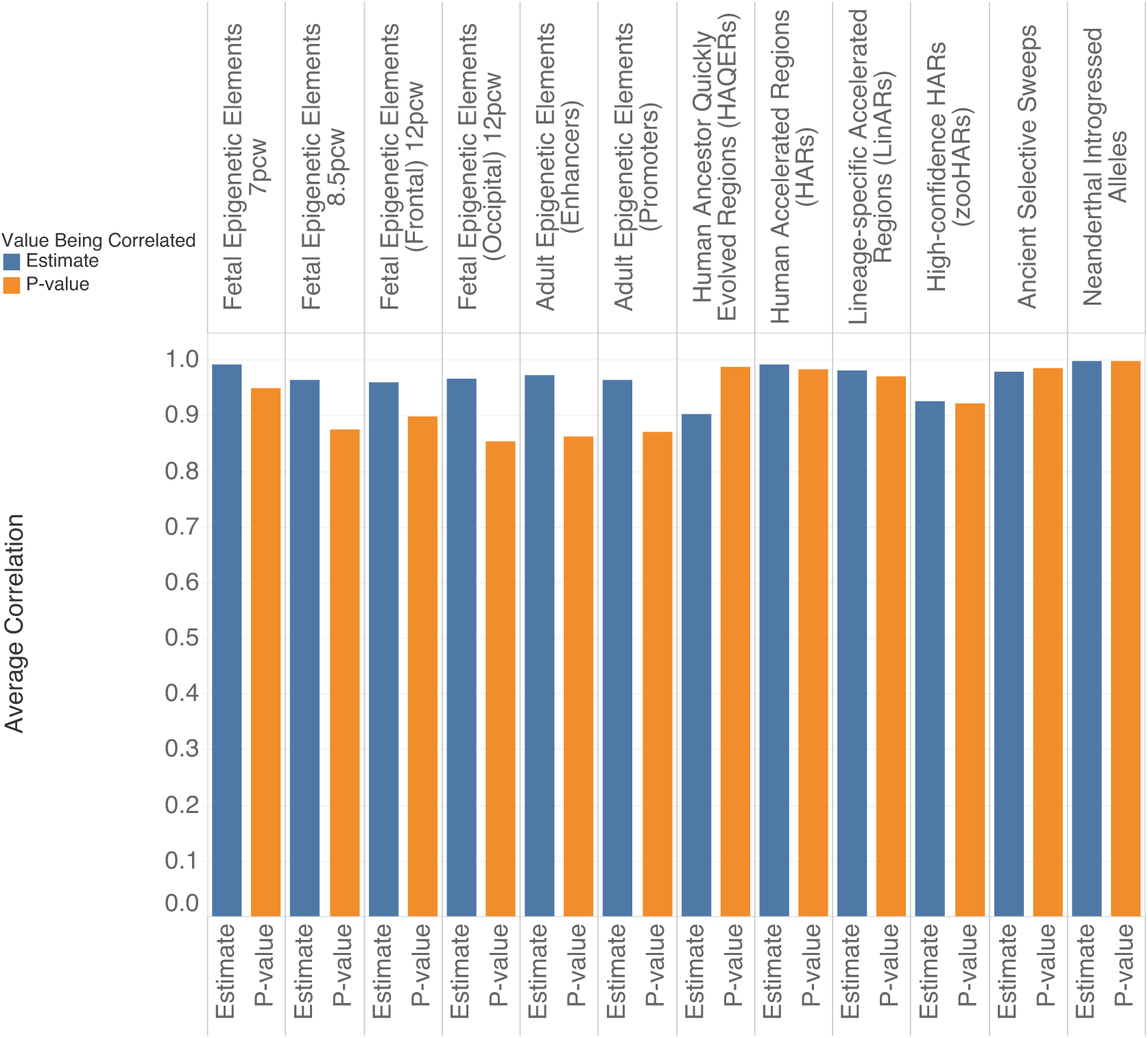
Comparison of annotation robustness. Average estimate and p-value correlations between enrichment analyses carried out on the full annotation set with three different subsets of 90% of the annotation. All fetal and adult epigenetic element sets were analyzed with S-LDSC while the remaining genomic regions were analyzed with HARE.

**Fig. S3.**
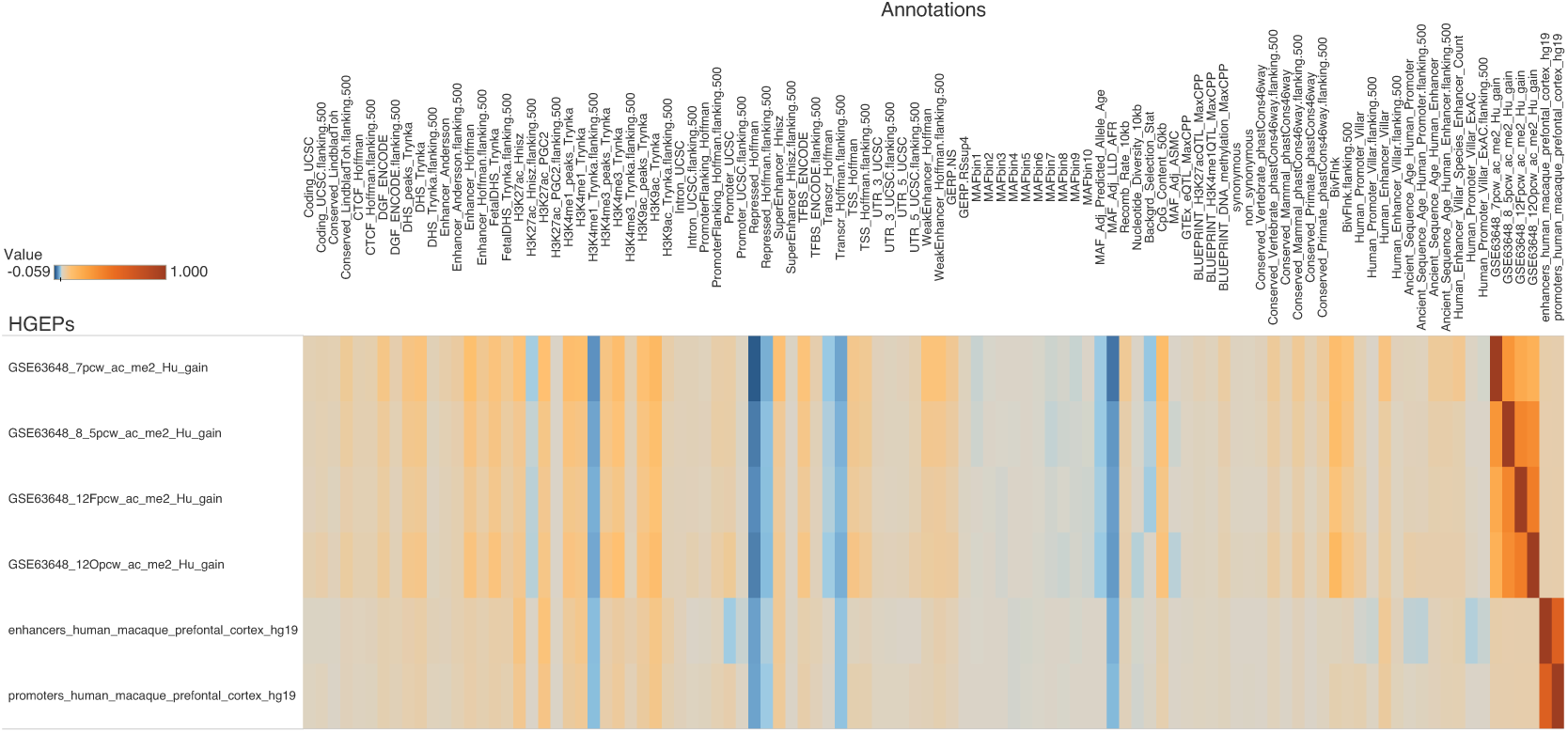
Correlations between baselineLDv2.2 annotations and HGEPs.

**Fig. S4.**
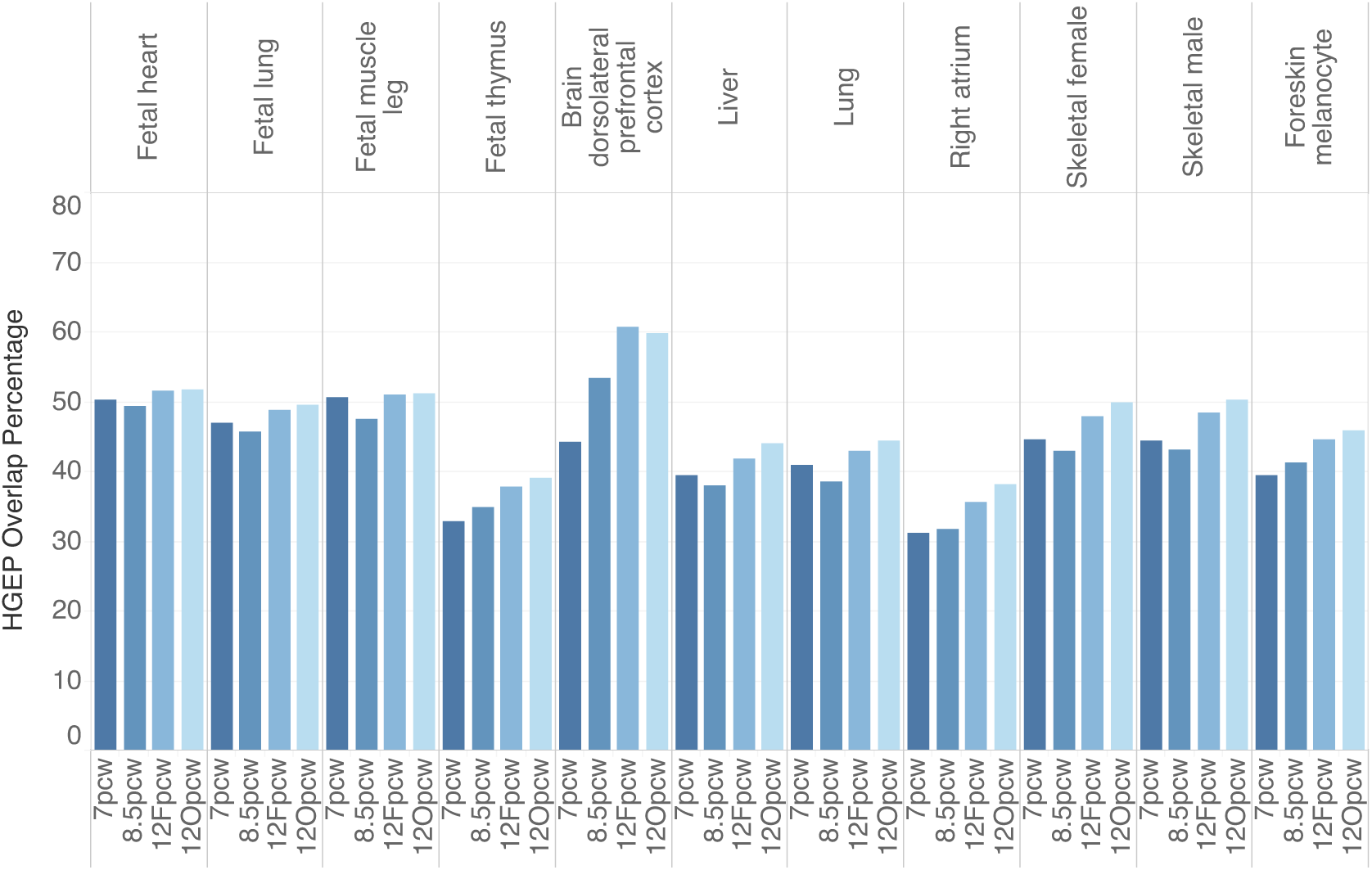
Comparison of base pair overlap. Shared overlap of base pairs between enhancers and promoters of various tissues with HGEP in the brain at various fetal time points. A total of 11 tissues obtained from various origins from the Epigenome Roadmap 25 state model were analyzed. The percent overlap is calculated as the number of bases that overlap between each enhancer/promoter annotation with the Epigenome Roadmap tissue divided by the total length of the enhancer/promoter annotation - i.e. (base pair overlap of 7 p.c.w and E066 (liver) / length of 7 p.c.w)

**Fig. S5.**
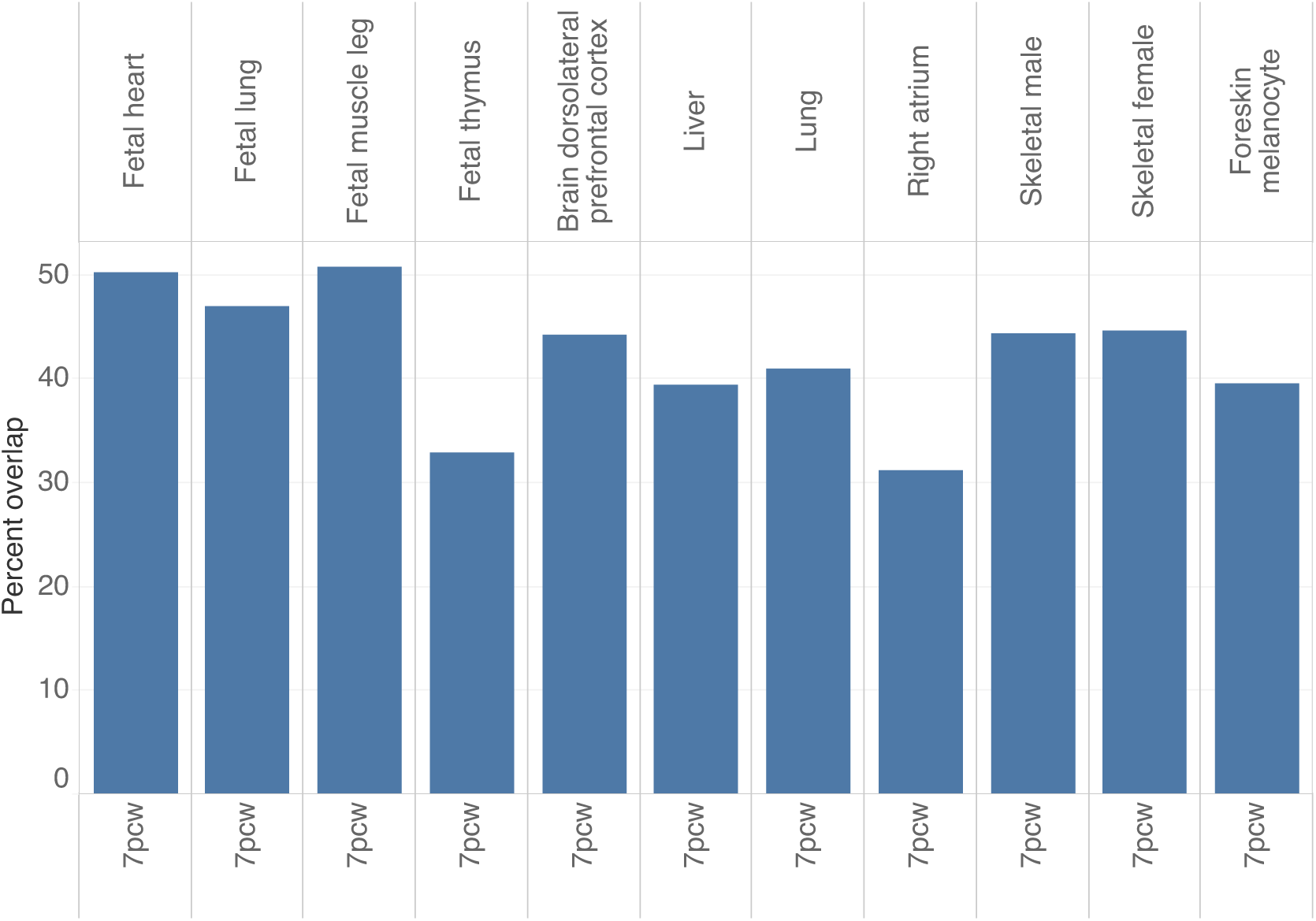
Shared overlap of base pairs between enhancers and promoters of various tissues with HGEP in the brain at 7 p.c.w only.

## Data and materials availability

The HARE enrichment pipeline is available at https://github.com/ossmith/HARE.

**Table S1** - GWAS phenotypes

**Table S2** - Genetic correlations

**Table S3** - Evolutionary Annotations

**Table S4** - S-LDSC Results

**Table S5** - S-LDSC Tissue Specificity

**Table S6** - HARE Results

**Table S7** - S-LDSC Meta-analysis

**Table S8** - HARE Meta-analysis

**Table S9** - HARE Robustness 1

**Table S10** - S-LDSC Robustness

**Table S11** - HARE Robustness 2

**Table S12** - Epigenome Roadmap Annotations

**Table S13** - Epigenome Roadmap State 25 Model

**Table S14** - Annotation Overlap

**Table S15** - HARE HGEP Results

## STAR Methods

### Key resources table

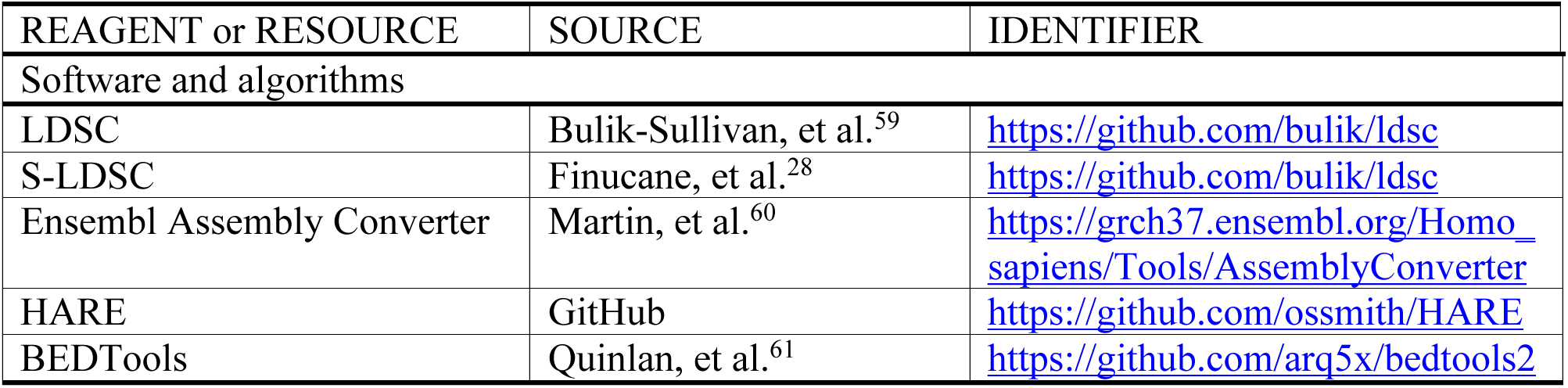

### Resource availability

#### Lead contact

Further information and requests for resources and reagents should be directed to and will be fulfilled by the lead contact, Eucharist Kun.

#### Materials availability

This study did not generate new unique reagents or materials.

#### Data and code availability

- This paper does not report original code, but all code used are listed in the key resources table and are publicly available as of the date of publication.
- Any additional information required to re-analyze the data reported in this paper is available from the lead contact upon request.

## Method details

### GWAS datasets

We analyzed a set of 41 independent GWAS summary statistics that were previously curated and analyzed^21,62^. These were curated by screening 34 GWAS summary statistics that are publicly available and 55 UK Biobank traits for which summary statistics were computed using BOLT-LMM (up to N= 459,324 European-ancestry samples). BOLT-LMM software is available at https://data.broadinstitute.org/alkesgroup/BOLT-LMM and the BOLT- LMM summary statistics for UK Biobank traits are available through the Alkes group. The set was first filtered to 47 traits

with z-scores of total SNP heritability of at least 6^62^. Since the set of 47 included 6 traits that were duplicated in two different datasets (genetic correlation >= 0.9), the final set of GWAS was reduced to 41 independent GWAS datasets (N=10,263 – 459,324; average N=320,000). We also analyzed GWAS obtained from 3 imaging studies carried out on UKB participants. We obtained 16 brain MRI traits (average N = 31,435), 10 right heart MRI traits (N = 41,135), and 6 skeletal DXA traits (N = 31,221)^11,32,33^. The brain MRI traits were chosen by taking the trait with the highest heritability from each MRI trait category as denoted by Smith et al^33^. Traits within each imaging study that had a genetic correlation higher than 0.9 were again filtered out. In total, we analyzed a set of 70 traits (**Table S1** and **Table S2**).

### Evolutionary genomic annotations

Genomic annotations were obtained from the following sources: (A) Epigenetic elements that gained novel function in the brain since our divergence with rhesus macaque at different developmental stages or post-conception weeks (p.c.w) and adulthood (Fetal human-gained (HG) enhancers and promoters at 7 p.c.w, 8.5 p.c.w, and 12 p.c.w and adult human-gained (HG) enhancers and promoters)^19,20^, (B) The fastest evolving regions of the human genome when compared to various sets of mammals and primates (human accelerated regions (HARs), lineage-specific accelerated regions (LinARs), high-confidence HARs (zooHARs), and human ancestor quickly evolved regions (HAQERs))^16,17,34,35^, (C) Ancient selective sweeps from the extended lineage sorting method capturing human-specific sweeps relative to Neanderthal/Denisovan^36^, and (D) Putatively introgressed variants from Neanderthals (**Table S3**)^37^.

### Annotation conversion to GRCH37

Annotations that were not in GRCH37 format were converted from GRCH38 to GRCH37 using the Ensembl Assembly Converter tool available at https://grch37.ensembl.org/Homo_sapiens/Tools/AssemblyConverter and stored in BED format. Resulting regions were merged using BEDTools ‘merge’^61^.

### Stratified LD score regression (S-LDSC) framework

Stratified LD score regression (S-LDSC) was developed in Finucane et al 2015^28^. The framework takes as input summary statistics from a GWAS, and LD-scores for SNPs in a set of test and control genomic categories. S-LDSC estimates whether a genomic region is enriched or depleted in heritability for a set of traits, capturing the contribution of variants in that genomic region towards trait variation, and whether this contribution is more or less than expected given the relative proportion of variants in that region. The per-SNP heritability of a SNP *i* and heritability enrichment of a genomic annotation or category are estimated from:

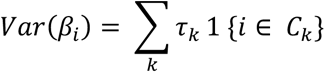

Where total heritability is represented as:

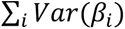

And heritability in category *C*_k_:

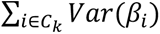

Then heritability enrichment (*h*^2^(*C*)) in category *C*_k_ is calculated as:

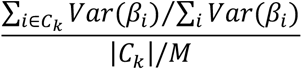

Lastly, jackknife standard error estimates generated by S-LDSC are used to compute confidence intervals for *h*^2^(*C*). Further details are outlined in Finucane et al 2015.

To account for multiple significance testing, the false discovery rate (FDR) was controlled at 0.05 using the approach of Benjamini-Hochberg^63^.

### S-LDSC genomic panels

The framework of S-LDSC uses three SNP panels. Here, we describe each and the set of SNPs used for each in our analysis, following previous analyses using S-LDSC^21,28,30^.

Reference SNPs (9,997,231 SNPs): These are SNPs used by S-LDSC to compute LD scores. We used the 1000 Genomes Phase 3 SNPs^64^ from the European super population with a minor allele count > 5.

Regression SNPs (1,187,349 SNPs): These are SNPs used by S-LDSC to estimate *τ_k_* from marginal association statistics. We used the HapMap Project Phase 3 SNPs (MHC region excluded)^65^.

Heritability SNPs (5,961,159 SNPs): These are SNPs used by S-LDSC to compute *h*^2^, *h*^2^(*C*), |*c*| and *sd_c_*. We used common variants (MAF > 0.05) among the set of reference SNPs (see above).

### S-LDSC joint model and control annotations

We used the S-LDSC framework to estimate the heritability enrichment in a given genomic annotation while jointly modeling the same parameters across a set of genomic annotations (Fig 3) (**Table S4**). The idea of the joint model is to account for confounding effects of other genomic annotations while estimating the heritability enrichment of a given annotation. The baseline model was presented in Finucane et al^28^ and includes a set of 53 annotations, those that partition the gene body as well as several epigenetic annotations. The baselineLDv1.1^30^ further includes a set of annotations related to selection such as allele frequency, allelic age, LD, nucleotide diversity, GERP scores and background selection for a total of 75 annotations. The BaselineLDv2.2^21,28–30^ is the most comprehensive and conservative set of annotations to include in a joint model with a test annotation and includes 97 genomic annotations (**Fig. S3**). This is the recommended baseline model and further includes annotations relating to conservation and age of epigenetic marks and whether a promoter is a promoter of an ExAC gene^21^. All heritability enrichment analyses in this study jointly model test annotations with the baselineLDv2.2 model. In our analysis of fetal human-gained regulatory elements, we further included in our model fetal brain regulatory elements from the Epigenome Roadmap 25 State Model^38^, taking the union of male (E081) and female (E082) marks. In our analysis of adult human-gained regulatory elements, we further included in our model adult brain prefrontal cortex regulatory elements (E073) from the Epigenome Roadmap 25 State Model^38^. For the 25 state model, a chromatin state model based on the imputed data for 12 marks, H3K4me1, H3K4me2, H3K4me3, H3K9ac, H3K27ac, H4K20me1, H3K79me2, H3K36me3, H3K9me3, H3K27me3, H2A.Z, and DNase, across all 127 reference epigenomes with 25-states was learned^38^. We included the following states that correspond to all regulatory elements (**Table S12** and **Table S13**).

### Tissue specificity analysis

As a proportion of the human-gained elements in the brain also overlap with regulatory elements in other tissues (**Table S14**), we performed additional analysis to determine the main driver of heritability enrichment across the different time points. Firstly, we examined whether the enrichment signals in the fetal gained enhancers and promoters at 7 p.c.w were driven by a greater number of base pair overlap with other tissues such as the heart, lung, or bone when compared to the other fetal timepoints. We obtained the genomic regions pertaining to regulatory elements from various tissues in the Epigenome Roadmap 25 state model to examine regions common to all tissues as well as each set of fetal HGEP (**Table S12** – **Table S14**)^38^. The number of base pair overlap between the brain and other tissues increases from 7 p.c.w to 12 p.c.w, ruling out increased overlap in regulatory regions at earlier time points rather than later time points as the main driving factor behind the increased heritability enrichment we observe at 7 p.c.w (**Fig. S4** and **Fig. S5**). To further elucidate the driving factor behind enrichment in the 7 p.c.w HGEPs, we split the annotation into two sets, regulatory elements that overlapped with genomic regions shared by all Epigenome Roadmap tissues and elements that had no overlap and performed S-LDSC on each set. We then correlated the heritability enrichment values of each subset with the original annotation. We found that the subset of brain HGEPs that overlapped regulatory elements in other tissues had an enrichment correlation of 0.91 with the original annotation. Similarly, the subset of brain HGEPs that shared no overlap with regulatory elements in other tissues had an enrichment correlation of 0.90 with the original results. Our results also show that heritability enrichment for some traits was only significant in the subset of brain HGEPs that overlapped regulatory elements in other tissues (p-value < 0.05) (**Fig. S1**). However, we see that the fetal brain-specific HGEPs at 7 p.c.w do contribute significant heritability enrichment across various trait domains (dermatological, skeletal, and respiratory) suggesting pleiotropic effects across the brain-body axis from our early evolution and life history.

### Gene enrichment analysis for evolutionary annotations

We scanned for elevated levels of intersections between genes containing genome-wide significant SNPs and our genomic annotations through a modified version of the method outlined in Xu et al^66^. For each phenotype, we first created annotations of protein coding regions that lie on our genome-wide significant SNPs using Ensembl’s GRCh37 Variant Effect Predictor version 105. We selected the closest protein coding feature within 5,000 base pairs up- or downstream of the SNP. Using biotype categorizations identified by VEP, these protein coding features were: (“protein_coding”, “IG_C_gene”, “IG_D_gene”, “IG_J_gene”, “IG_LV_gene”, “IG_M_gene”, “IG_V_gene”, “IG_Z_gene”, “nonsense_mediated_decay”, “nontranslating_CDS”, “non_stop_decay”, “polymorphic_pseudogene”, “TR_C_gene”, “TR_D_gene”, “TR_J_gene”). We refer to the list of features for all independent genome-wide significant loci significantly associated with the trait as the *element set* for the phenotype being analyzed. Phenotypes with fewer than 50 elements in their set were removed from the analysis due to insufficient power. We then used BioMart command line queries to generate the genomic locations (chromosome, start, stop) of each feature within the human genome. In order to scan for selection, we used BEDTools ‘intersect’ to compute the number of intersections found in the gene set with genomic annotations sourced from literature.

To generate a background distribution of intersections per bp, we computed the annotation- element intersections per bp of 1,000 length-matched element sets. Because the distribution of these feature lengths is non-normal, we binned the element sets into deciles based on gene length and computed the average length *l* within each bin of size *n*. For each bin in the simulation, we sampled *n* random elements of length *l* to create our complete element set which was then used to compute the intersections per base pair of the simulated set. Due to the large differences in element set sizes and lengths across phenotypes, a background distribution was generated independently for each phenotype analyzed. On this background we fit a Weibull distribution for computation of p-values of the observed intersections in comparison to the background. Overlap enrichment was then calculated as a percent difference between GWAS gene overlap with an annotation compared to random gene overlap with an annotation. This was calculated by subtracting the number of intersections between genes related to a GWAS trait and the annotation of interest from the average number of intersections between random genes and the annotation of interest and then dividing by the average number of intersections between random genes and the annotation of interest^27^.

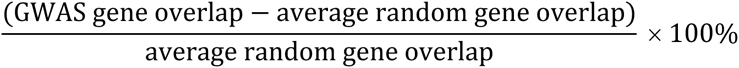

### S-LDSC and HARE meta-analysis

To meta-analyze results while accounting for genetic correlations among the various traits, we performed a random effects meta-analysis by each trait category using the rma.mv function from the metafor package. This computes a summary estimate and summary standard error from a collection of effect estimates and standard errors for each trait and also considers a genetic variance-covariance matrix among the traits. These were used to compute confidence intervals and z-scores to test for significance of the summary estimate for each annotation. To account for multiple significance testing, the false discovery rate (FDR) was controlled at 0.05 using the approach of Benjamini-Hochberg.

### Annotation robustness analysis

In order to assess the sensitivity and specificity of the various approaches used in finding these various evolutionary genomic annotations, we examined the impact of randomly sampling just 90% of each of the regions over 3 replicates and running S-LDSC and HARE on these subsets. We then correlated the corresponding enrichment estimates and p-values to the original results and found that these subsets had an average Pearson correlation of 95% with the original annotations (**Fig. S2**). Another approach we took was to shorten or lengthen the ends of every genomic region in each annotation by 5% and re-run HARE on the altered set of regions. The correlation between these sets and the original sets was greater than 99% (**Table S9** – **Table S11**). All correlations were performed with base R functions.

### Annotation comparison

Annotations were tested for base pair overlap using BEDTools^61^. Annotations were paired and the overlapping regions between the annotations was calculated using BEDTools ‘intersect’. The proportion of intersecting base pairs between two annotations with the length of each original annotation was then calculated (**Table S14**).

### Differences among various types of human accelerated regions

As the comparative genomic annotations were generated through slight differences in either the number or type of species used in genome alignment, we wanted to probe the variation across the 4 annotations. The set of HARs we examined were taken from a study that combined 4 datasets that used a variety of filters and limited multi-species alignment (generally chimpanzee, mouse, and rat genomes)^34^ while the LinARs and zooHARs were generated from aligning new long read sequences of many more species. In particular, LinARs were generated from aligning 49 high-quality reference genomes from primate species while the zooHARs were generated from an alignment of 241 mammalian genomes^16,17^. These 3 comparative genomic analyses focused on conserved regions of the genome, which is only 5% of the total human genome. Our last genomic annotation, HAQERs, were generated by performing comparative genomic analysis on only great ape species, which allowed for the discovery of fast evolving regions of the human genome present in the non-conserved region of the genome^35^. Comparison of the location and size of these various genomic annotations shows that HARs, LinARs, and zooHARs are closely related and share some amount of overlap while HAQERs are vastly different (**Table S14**). We found that 18% of LinARs overlapped HARs while 45% of zooHARs overlapped HARs.

However, in terms of gene enrichment, LinARs and HARs had an enrichment correlation of 0.71 while zooHARs and HARs only had an enrichment correlation of 0.42.

### S-LDSC and HARE comparison

We carried out HARE analysis on the fetal and adult human gained enhancers and promoters for all traits to see if the results were in agreement with S-LDSC results. Overall, the correlation between heritability enrichment and gene enrichment was low (Pearson correlation = 0.23), however traits that were significantly enriched in S-LDSC were also significant in HARE (FDR-adjusted p-value < 0.05) (**Table S15**).

